# Ion-Pair-Free Capillary HILIC-MS for Sensitive Nucleic Acid Analysis and RNA Modification Mapping

**DOI:** 10.64898/2026.08.19.745671

**Authors:** Junzhou Wu, Rana Togay, Jingjing Sun, Dwijapriya, Chi-Kong Chan, Alex Reading, Liang Cui, Xueming Dong, Peter Dedon

## Abstract

Mass spectrometry (MS)-based nucleic acid analysis provides direct chemical evidence for oligonucleotide sequence, composition, and modifications. However, oligonucleotide LC-MS analysis commonly relies on ion-pairing reversed-phase liquid chromatography (IP-RPLC). Although IP-RPLC provides strong retention and high-resolution separation of highly charged nucleic acids, ion-pairing reagents can contaminate LC-MS systems, suppress electrospray ionization, require extensive system cleaning, and limit the use of high-end MS platforms that are primarily dedicated to proteomics or metabolomics. Here, we developed and evaluated an ion-pair-free capillary hydrophilic interaction liquid chromatography mass spectrometry (capillary HILIC-MS) workflow for RNA modification mapping. To enable robust analysis of biologically relevant samples, we optimized sample preparation, high-organic loading conditions, chromatographic parameters, and MS source settings to overcome key challenges associated with capillary HILIC, including limited sample volume, solvent compatibility, and solvent breakthrough during injection. The optimized capillary HILIC-MS method provided effective separation of oligonucleotides below 30 nt and enabled sensitive detection of RNA modifications in the populations of tRNAs and rRNAs in biological samples. Importantly, the ion-pair-free workflow also allowed switching between nucleic acid analysis and proteomics on the same LC-MS platform without the need for extensive system decontamination. Together, this workflow provides a sensitive, robust, and MS-compatible approach for nucleic acid analysis, expanding the utility of high-end LC-MS systems for both therapeutic oligonucleotide characterization and biological RNA modification profiling.

## INTRODUCTION

DNA and RNA undergo extensive chemical modification catalyzed by hundreds of dedicated modifying enzymes. These modifications expand the functional capacity of the genome and transcriptome beyond their primary sequences, providing an additional layer of information regulating gene expression. In DNA, chemical modifications contribute to chromatin organization, transcriptional regulation, genome stability, and cellular identity [1-3]. In RNA, modifications regulate RNA processing, stability, localization, translation efficiency, and stress responses [4-7]. Because these processes are fundamental to normal cellular function, dysregulation of nucleic acid modifications has been closely linked to aging, cancer, developmental and metabolic disorders, neurological diseases, and host–pathogen interactions [3,4,6-11]. In addition to naturally occurring modifications, chemically modified nucleic acids are increasingly used in therapeutic platforms, including antisense oligonucleotides, siRNA, mRNA vaccines, and other RNA- or DNA-based therapeutics, where modifications are critical for stability, delivery, potency, and immunogenicity [12-15]. Therefore, accurate characterization of nucleic acid modifications is important not only for understanding biological regulation and disease mechanisms, but also for the development and quality control of nucleic acid therapeutics.

A variety of technologies has been developed to detect and characterize nucleic acid modifications [16,17]. Antibody-based methods, such as immunoblotting, ELISA, immunoprecipitation, and immunostaining, are relatively simple and scalable, but their specificity can be limited by antibody cross-reactivity, and they can’t provide positional information [17,18]. Chemical derivatization- and enzyme-based next-generation sequencing methods can enable transcriptome-wide mapping of selected RNA modifications but are generally limited to specific modification types [17,19-21]. Nanopore-based approaches have further expanded the ability to detect multiple modifications, but data interpretation can be challenging and often requires extensive optimization, training datasets, or orthogonal validation [22-24]. In contrast, liquid chromatography–mass spectrometry (LC-MS) provides direct chemical measurement of modified RNA fragments with high sensitivity, specificity, and quantitative accuracy [25-30]. However, MS-based approaches also face challenges, including sample preparation complexity, incomplete sequence coverage, limited sensitivity in complex biological samples, and a strong dependence on chromatographic strategies that often require ion-pairing reagents, which can compromise MS sensitivity, robustness, and instrument compatibility.

Ion-pair reversed-phase (IP-RP) LC-MS has been widely used for nucleic acid analysis because ion-pairing reagents improve chromatographic retention and separation of highly negative charged nucleic acid sequences [31]. However, these reagents can suppress electrospray ionization, contaminate LC-MS systems, and require extensive equilibration and cleaning [32,33]. Thus, a dedicated IP-RP LC-MS system is typically required for nucleic acid analysis. This limits nucleic acid analysis in shared core facilities and industrial settings, particularly on high-end LC–MS systems that are also routinely used for proteomics or metabolomics. In recent years, hydrophilic interaction liquid chromatography (HILIC) has emerged as an attractive ion-pair-free alternative for separating polar nucleosides and oligonucleotides using MS-compatible volatile buffers [34-36]. This trend is also reflected in industry efforts to develop bio-inert HILIC analytical columns for nucleic acid analysis, such as the Altura oligo HILIC column from Agilent Technologies and the Premier BEH amide column from Waters. Further miniaturization to capillary- and nano-flow HILIC-MS offers significantly increased sensitivity, which is particularly valuable for low-input biological samples and low-abundance nucleic acid modifications. However, capillary- and nanoflow HILIC presents its own technical challenges. Because HILIC requires samples to be loaded in a high-organic solvent, excessive aqueous content can cause solvent breakthrough, peak distortion, reduced analyte retention, and substantial loss of sensitivity [37,38]. A recently reported ion-pair-free nanoflow HILIC-MS workflow enabled analysis of 25–50 ng of a single purified yeast tRNA species [39], highlighting the potential of nanoflow HILIC for low-input RNA analysis. Nevertheless, the broader application of capillary- and nanoflow HILIC–MS to complex biological RNA mixtures, particularly when sample availability is limited, remains relatively underexplored.

Here, we present an ion-pair-free capillary HILIC-MS workflow that has been successfully applied to the analysis of biological RNA samples. This workflow incorporates a practical sample preparation protocol that enables limited biological samples to be prepared in small volumes of high-organic loading buffer, together with carefully optimized analytical conditions, including buffer composition, chromatographic parameters, and MS source settings. We evaluate the performance of the workflow in triplicate analyses of HEK293 rRNA and PA14 tRNA samples. Based on modifications reproducibly detected across the three replicate injections, 8 modification types were mapped to 56 sites in HEK293 rRNA, including nine partially modified sites, while 22 modification types were mapped to 68 sites in PA14 tRNA. Importantly, the LC-MS system remained compatible with subsequent proteomics analysis, with no need for extensive system cleaning. Together, this workflow improves compatibility with high-end LC-MS platforms and provides a sensitive, reproducible, and MS-friendly approach for nucleic acid modification analysis in low-input and complex biological samples.

## MATERIALS AND METHODS

### Preparation of poly(T) standard mixtures

The poly(T) oligonucleotide standard mixture (10 μM) was prepared by combining equimolar amounts of desalted oligonucleotides containing 5, 10, 15, 20, 25, or 30 thymidine residues (GenScript Biotech). For analyses using the analytical column and selected capillary-column experiments, each oligonucleotide was prepared at a final concentration of 1 μM. For the remaining capillary-column analyses, each oligonucleotide was prepared at 0.2 μM. In all cases, samples were prepared in 70% acetonitrile containing 10 mM ammonium acetate.

### HEK293 cell culture, rRNA isolation, and sample preparation

HEK293 cells were cultured using DMEM supplemented with 10% FBS and 1% Penicillin-Streptomycin at 37 °C in a humidified incubator containing 5% CO_2_. Cells were harvested at approximately 80% confluence. The large-RNA fraction, enriched in 18S and 28S rRNAs, was isolated using the PureLink miRNA Isolation Kit (Thermo Fisher Scientific, K157001) according to the manufacturer’s instructions and recovered from the first spin cartridge. Purified rRNA (10 µg) was digested with 1,000 U of RNase T1 (Thermo Fisher Scientific, EN0541) in 50 µL of 1× TE buffer at 37 °C for 30 min. Following digestion, 50 µL of phenol:chloroform:isoamyl alcohol (25:24:1, pH 6.6; Ambion, AM9730) was added. The samples were vortexed and centrifuged to separate the aqueous and organic phases. The aqueous phase was carefully collected without disturbing the interphase and transferred to a new tube. RNA fragments were recovered using the Oligo Clean & Concentrator Kit (Zymo Research, D4060). Briefly, 100 µL of binding buffer and 800 µL of ethanol were added to each sample, and purification was performed according to the manufacturer’s instructions. RNA fragments were eluted in 18 µL of water, followed by the addition of 2 µL of 1 M triethylammonium acetate (TEAA; Honeywell Fluka, 90357).

The samples were further purified using Agilent OMIX C18 10 µL pipette tips (Agilent Technologies, A5700310). The tip was prewetted twice with 10 µL of 50% acetonitrile in water and equilibrated by three washes with 10 µL of 0.1 M TEAA. The sample was loaded by aspirating and dispensing it through the tip 5–10 times. The tip was then washed three times with 10 µL of fresh 0.1 M TEAA, followed by three washes with 10 µL of water. The final samples were eluted in 8 µL of 70% acetonitrile containing 5 mM ammonium acetate and used directly for LC–MS/MS analysis.

### PA14 culture, small RNA isolation, and sample preparation

*Pseudomonas aeruginosa* UCBPP-PA14 wild-type was obtained from the non-redundant PA14 transposon insertion mutant library [40]. The strain was initially cultured overnight at 37 °C with shaking at 300 rpm in 2 mL of LB medium in a 5 mL round-bottom tube. Subsequently, 100 µL of the overnight culture was transferred to 5 mL of fresh LB medium in a 14 mL round-bottom tube and incubated at 37 °C with shaking at 300 rpm until the culture reached an OD_600_ of ∼0.8. Cells were then pelleted by centrifugation of 3,000 × g for 10 min at 4 °C. Medium was removed and cells were washed with cold 1× PBS buffer. Cell lysis buffer (100 µL; 50 mM Tris-HCl, 1 mM EDTA, 4 M guanidine isothiocynate, pH 7.5) was added and the plate was shaken vigorously at ambient temperature (1500 rpm) for 10 min to ensure complete cell lysis.

Small RNA (∼80-90% tRNA) was isolated using an in-house magnetic bead–based protocol [9]. Briefly, 45 µL of cell lysate was mixed with 105 µL of RNA binding buffer I containing 3 M LiCl, 7% PEG8000, 2 mM EDTA, 40 mM Tris (pH 7.5), and 8 µL Sera-Mag carboxylate-modified magnetic beads (Cytiva, 65152105050350). The mixture was agitated thoroughly and incubated at ambient temperature for 5 min. The magnetic beads were separated on a magnetic rack at ambient temperature for 5 min and the supernatant was transferred to a new tube. The tube was then centrifuged at 3000 xg for 1 min and placed on the magnetic rack to remove any remaining beads. The supernatant was transferred to a new tube and mixed with 1.8 x (v/v) RNA-binding buffer II (0.05% magnetic beads in isopropanol). The mixture was agitated thoroughly at ambient temperature for 5 min. The magnetic beads were separated on the magnetic rack for 5 min and the supernatant was removed. The pelleted beads were washed twice using washing buffer (10 mM Tris-HCl, 80% EtOH, pH 7.5). The beads were air dried at ambient temperature. tRNA was eluted in 50 µL nuclease-free water and stored at −20 °C until further processing. Subsequent RNase T1 digestion and sample preparation were performed as described above for HEK293 rRNA.

### LC-MS/MS data acquisition using capillary HILIC column

Capillary HILIC separations were performed on an Easy-nLC 1000 system (Thermo Fisher Scientific). The capillary HILIC column was packed using the BEH amide stationary phase (130Å, 1.7 µm) obtained from unpacking a Waters Acquity UPLC BEH amide column. The column was prepared using a 25 cm length of fused-silica tubing (150 μm ID, 360 μm OD; IDEX, FS-115) fitted with a porous silica frit at one end, prepared using a frit kit (Next Advance). The column was packed in a pressure injection cell (Next Advance) using a slurry of BEH Amide stationary-phase particles in acetonitrile until a packed-bed length of 13 cm was achieved. The packed column was trimmed to the length of packed-bed, connected to an EASY-Spray capillary emitter and coupled to an Orbitrap Exploris 240 mass spectrometer (Thermo Scientific, USA). Chromatographic separation was carried out at a constant flow rate of 1200 nL/min. Mobile phase A consisted of 80% ACN (80:20 v/v ACN:Milli-Q water) with 5 mM AmAA, and mobile phase B consisted of 40% ACN (40:60 v/v ACN:Milli-Q water) with 5 mM AmAA. For Poly(T) standards, the gradient start from 2% B and inceased to 30% over 1 min, then increased from 30% to 60% B over 5 min, to 75% B at 10 min and 85% B at 15 min, reached 100% B at 16 min, was held until 20 min, and returned to 2% B by 22 min. For biological samples, the gradient began at 2% B, increased to 40% B over 1 min, to 70% B at 16 min and 90% B at 31 min, reached 100% B at 46 min, was held for 4 min, and returned to 2% B by 52 min, with re-equilibration to 55 min.

Mass spectrometric analysis was performed in negative ion mode using MS1 for poly(T) standards and data-dependent acquisition (DDA) MS/MS for biological samples. The source voltage was set to 1.4 kV. The ion transfer tubing temperature was set to 350 °C. Full MS scans were acquired over an *m/z* range of 570–2000 at a resolution of 60,000, with an AGC target of 300% and a maximum injection time of 100 ms. Each full MS scan was followed by 3 s of MS/MS acquisition, and dynamic exclusion was enabled after one selection for 15 s. MS/MS spectra were acquired over an *m/z* range of 110–2000 at a resolution of 60,000, with an AGC target of 1000% and a maximum injection time of 500 ms. Higher-energy collisional dissociation (HCD) was performed using normalized collision energies of 24, 27, and 30. The precursor isolation window was set to 2 Da.

### LC-MS/MS data acquisition using analytical HILIC column

Analytical-flow HILIC separations were performed using a Vanquish UHPLC system (Thermo Fisher Scientific) equipped with an ACQUITY UPLC BEH Amide column (2.1 × 150 mm, 130 Å, 1.7 µm; Waters). The column temperature was maintained at 60 °C, and chromatographic separation was performed at a constant flow rate of 0.30 mL/min. Mobile phase A consisted of 80% acetonitrile in Milli-Q water (80:20, v/v) containing 10 mM ammonium acetate (AmAA), whereas mobile phase B consisted of 40% acetonitrile in Milli-Q water (40:60, v/v) containing 10 mM AmAA. The gradient began at 0% B, increased to 30% B at 1 min, 55% B at 10 min, and 70% B at 20 min, and reached 100% B at 21 min. The gradient was maintained at 100% B until 23 min, returned to 0% B at 24 min, and was held at 0% B until 30 min for column re-equilibration. The UHPLC system was coupled to an Orbitrap Exploris 240 mass spectrometer equipped with a heated electrospray ionization source. Data were acquired in negative-ion mode using a spray voltage of 2.5 kV. The sheath-gas and auxiliary-gas flow rates were set to 50 and 10 arbitrary units, respectively. The ion transfer tubing and vaporizer temperatures were set to 325 °C and 350 °C, respectively. Full-MS acquisition parameters were identical to those used for the capillary HILIC–MS analysis.

### OpenMS-based oligonucleotide identification and feature detection

Raw data acquired from the Orbitrap mass spectrometer were converted to mzML format using the MSConvert GUI tool (Version: 3.0.25239-8e11d5a) from ProteoWizard [41]. mzML data were then processed using an OpenMS (version 3.5.0) workflow [29] in TOPPAS. A target/decoy sequence database was generated from the input RNA FASTA file using *DecoyDatabase*, with the sequence type set to RNA and decoy sequences generated by shuffling. Each mzML file was searched independently against the target/decoy database using *NucleicAcidSearchEngine*. RNase T1 was specified as the digestion enzyme, with up to one missed cleavage allowed and a minimum oligonucleotide length of 5 nt. The precursor and fragment-ion mass tolerances were both set to 20 ppm. A maximum of two variable modifications was permitted per oligonucleotide, and identifications were filtered using a false-discovery-rate cutoff of 0.1. For label-free feature detection, the target list generated by *NucleicAcidSearchEngine* was supplied to *FeatureFinderMetaboIdent* together with the corresponding mzML file. This step extracted and integrated MS1 chromatographic features associated with the identified oligonucleotides, generating feature-level outputs for subsequent quantitative analysis. Finally, each mzML file was opened in TOPPView, and the corresponding idXML output was loaded using *Tools > Annotate with peptide identifications.* The annotated results were then exported from TOPPView as a TSV file containing the identified oligonucleotide fragments in tabular format.

### Downstream data analysis and visualization

Downstream data analysis was performed using custom Python scripts. RNA fragment identifications from three replicate injections were consolidated by sequence, and fragments detected in all three injections were retained for reproducibility and sequence-coverage analyses. Within each injection, signal intensities were summed for each sequence, normalized to the total identified signal, and log₁₀-transformed. Intensity reproducibility was evaluated through pairwise comparisons of normalized feature peak areas and precursor-ion intensities.

OpenMS featureXML files were parsed to extract retention time, peak area, charge state, precursor *m/z*, and peak-model status for each feature. Features were matched across injections according to charge state and precursor *m/z*, using a mass tolerance of 5 ppm. Retention-time reproducibility was assessed by pairwise Pearson correlation analysis. Features assigned a peak-model status of 0, indicating a valid fitted peak model, in both injections were distinguished from features with missing or non-valid peak models.

Sequence coverage was mapped to the human 18S and 28S rRNA reference sequences. Fragments that mapped to multiple positions were retained at all compatible locations. Experimentally detected fragments, manually curated identifications, and theoretical RNase T1 digestion products of at least 6 nt were converted into contiguous coverage intervals for visualization. Unlike the OpenMS search, this *in silico* RNase T1 digestion incorporated reference modification annotations, with cleavage after guanosine suppressed at annotated Gm and m^7^G residues.

## RESULTS AND DISCUSSION

### Development and optimization of the capillary HILIC-MS platform

The hardware setup for capillary HILIC-MS was not previously well established. Capillary HILIC columns are not widely available, and several commercially available options still use stainless-steel hardware, which is not ideal for nucleic acid analysis. Thus, we decided to make self-packed column using 150 µm ID fused-silica tubing with a single-end porous silica frit and packed using a pressure injection cell. The stationary phase was BEH amide particles (1.7 µm, 130 Å; Waters Corporation). The column was connected to an EASY-Spray capillary emitter with a 15 µm ID tip (Thermo Fisher Scientific). This hardware configuration established a practical bioinert capillary HILIC-MS platform with a metal-free flow path suitable for nucleic acid analysis (**Fig. 1a**).

**Figure 1.**
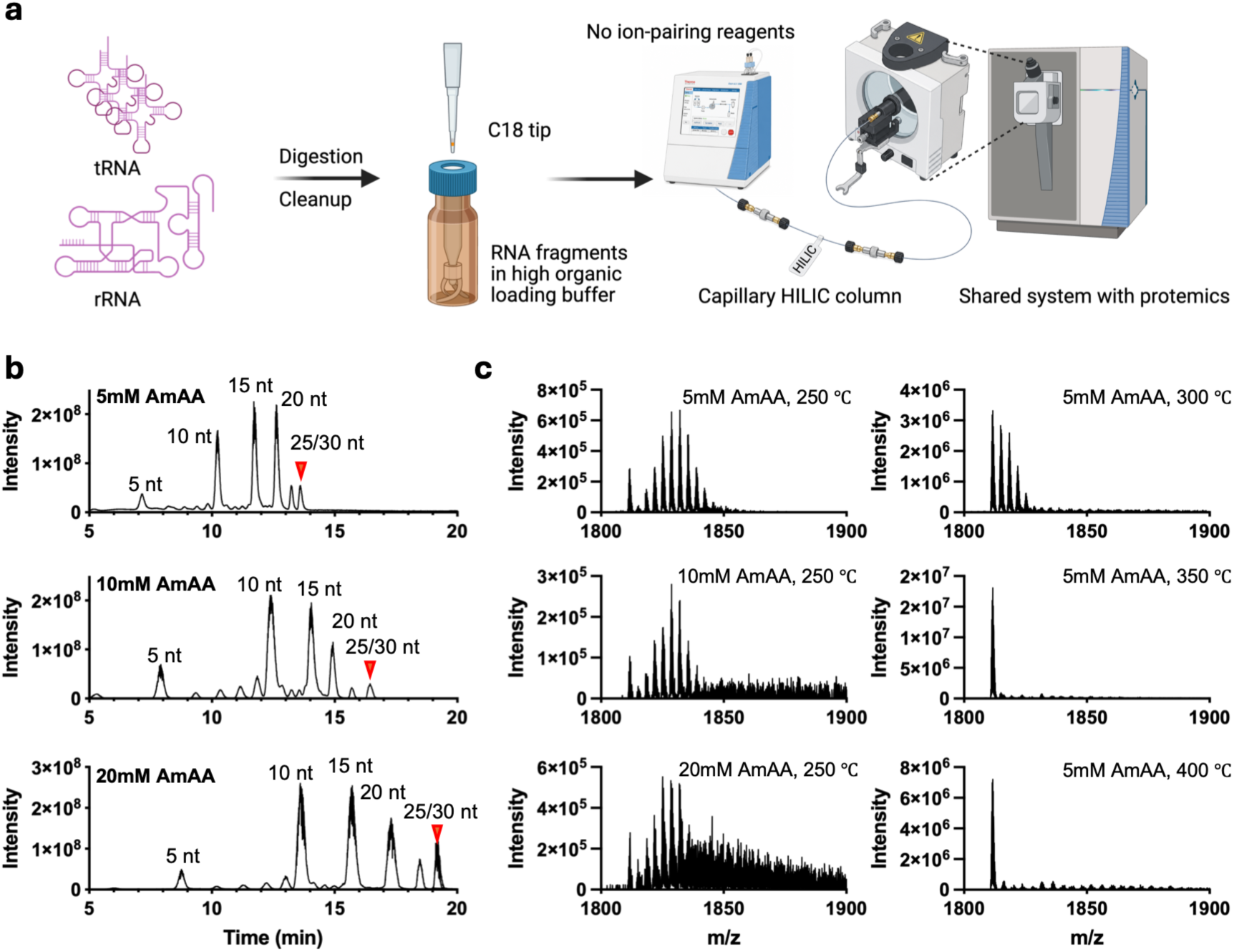
Optimization of LC–MS conditions for capillary HILIC–MS analysis. (**a**) Overview of the sample-preparation workflow and instrumental setup for capillary HILIC–MS analysis. (**b**) Total ion chromatograms (TICs) of a synthetic poly(T) ladder acquired using different ammonium acetate (AmAA) concentrations. (**c**) MS1 spectra of the 30-nt poly(T) oligonucleotide (z = −5) acquired using different AmAA concentrations and ion-transfer tube temperatures.

Because aqueous sample loading was incompatible with capillary HILIC columns, which have a column volume of only approximately 2 µL, we first evaluated loading buffers containing 50–80% acetonitrile (ACN) with 5 or 10 mM ammonium acetate (AmAA) using a series of poly(dT) oligonucleotides ranging from 5 to 30 nt (**Fig. S1**). A sample volume of 1 µL was injected for all analyses. At 50% ACN, the 5-, 10-, and 15-nt oligonucleotides produced broad, small or split peaks, consistent with solvent breakthrough during sample loading. At 60% ACN, the 5-nt oligonucleotide continued to show a split peak, whereas the longer oligonucleotides exhibited improved peak shapes. At 70% ACN, all oligonucleotides produced well-defined chromatographic peaks at both 5 and 10 mM AmAA buffers. Increasing the ACN concentration to 80%, however, resulted in the loss of the longer oligonucleotides. This effect was not observed when the same sample was analysed using the analytical-scale system, suggesting that oligonucleotide loss may have occurred within the flow path of the EASY-nLC system instead of percipatation in loading buffer. Collectively, these results identified 70% ACN containing 5 or 10 mM AmAA as the optimal loading buffer, providing effective retention across the tested oligonucleotide lengths while minimizing solvent breakthrough and precipitation.

Further optimizations focused on the AmAA concentration in the gradient mobile phases and the ion transfer tube temperature. We first compared 5, 10, and 20 mM AmAA under the same gradient conditions. At all three conditions, the capillary HILIC column provided effective separation of poly(T) oligonucleotides ranging from 5 to 30 nt (Fig. 1b). Increasing the AmAA concentration improved oligonucleotide retention but also promoted ammonium adducts formation, particularly for longer oligonucleotides (Fig. 1c). For the 30 nt oligonucleotides, no clear molecular ion peak was observed under the 20 mM AmAA condition due to extensive adduct formation. To reduce ammonium adducts, the ion transfer tube temperature was then increased stepwise from 250 °C to 400 °C. At temperatures of 350 °C and above, ammonium adducts were substantially reduced, and the MS1 spectrum showed a clear molecular ion peak (**Fig. 1c**). Based on these results, we established the final capillary HILIC-MS conditions using 5 mM AmAA in the gradient mobile phases, and an ion transfer tube temperature of 350 °C for further analysis.

### Performance comparison between analytical-flow and capillary-flow HILIC-MS

We further compared the chromatographic and MS performance of analytical and capillary HILIC columns using poly(T) oligonucleotides mixture. The analytical column provided good chromatographic resolution and peak shape across the tested size range, with peak widths at half height of 0.05–0.08 min. The capillary HILIC column also achieved reasonable separation, although broader peaks were observed, with peak widths at half height ranging from 0.08 to 0.32 min (**Fig. 2a**). Retention times were highly stable across repeated injections on the analytical columns (**Fig. S3**). In comparison, the capillary HILIC column showed minor retention-time shifts of less than 0.06 min. The capillary setup also exhibited lower MS signal stability, as reflected by greater variability in the TICs, than the analytical-column system equipped with a conventional ESI source (**Fig. 1b, 2a**).

**Figure 2.**
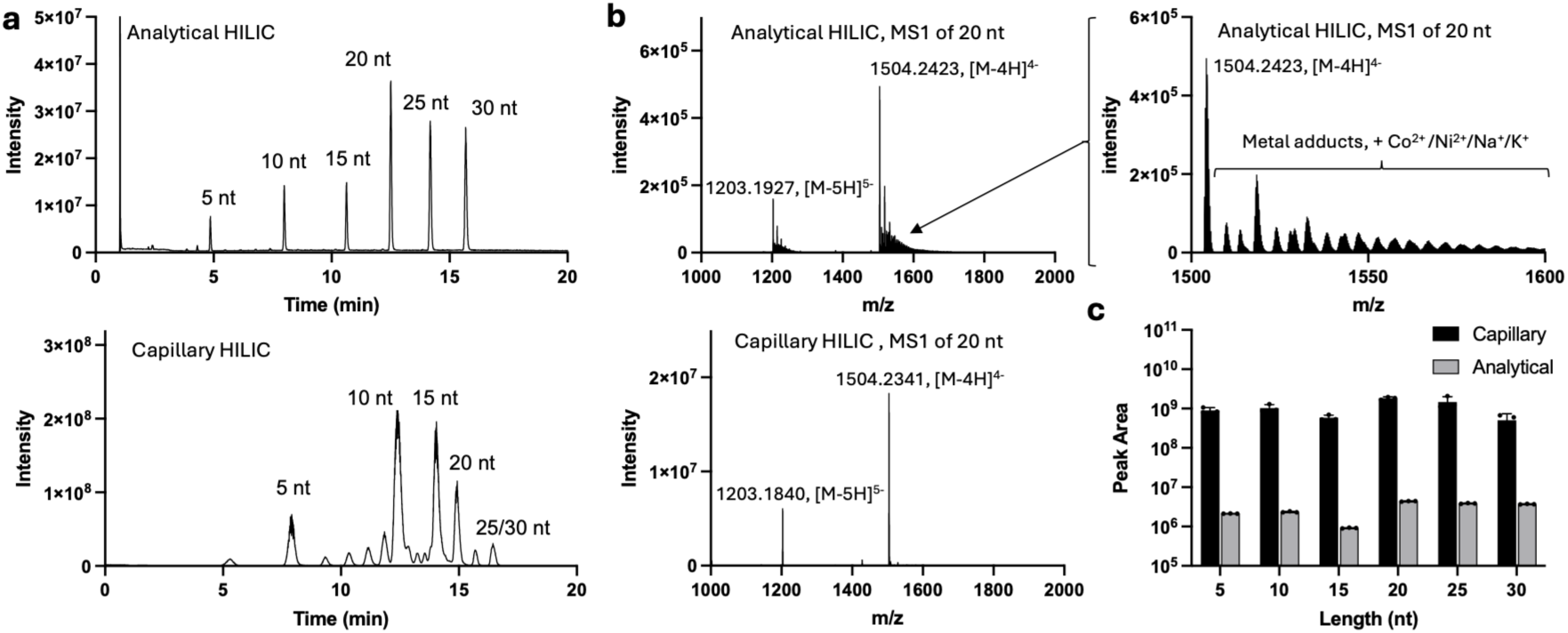
Comparison of analytical- and capillary-scale HILIC–MS. (**a**) TICs of a synthetic poly(T) ladder analyzed by analytical HILIC–MS (top) and capillary HILIC–MS (bottom). (**b**) MS1 spectra of the 20-nt poly(T) oligonucleotide acquired using analytical HILIC–MS (top) and capillary HILIC–MS (bottom). The expanded m/z 1500–1600 region (right) highlights ion adducts observed in the analytical HILIC–MS spectrum. (**c**) Comparison of peak areas obtained from 1-pmol injections of the poly(T) ladder using analytical- and capillary-scale HILIC–MS.

Despite its reduced chromatographic performance, the capillary HILIC-MS setup produced significant improvements in MS sensitivity, background reduction, and adduct suppression. As an example, the MS1 spectrum of the 20-nt poly(T) oligonucleotide obtained using the capillary column showed clear −4 and −5 charge states with minimal adduct formation. In contrast, the analytical HILIC system produced prominent metal-adduct peaks that were not limited to the commonly observed Na^+^ and K^+^ adducts. Accurate-mass measurements and isotope-envelope analysis indicated that the major transition-metal adducts were consistent with a mixture of Ni(II)- and Co(II)-containing species (**Fig. S3**). These adducts may have originated from MP35N alloy-containing components in the LC flow path, which are commonly used in bioinert LC systems. Overall, the capillary platform showed 130- to 640-fold increase in peak area compared with the analytical LC-MS setup, depending on oligonucleotide length (**Fig. 2c**). These results demonstrate that miniaturized capillary HILIC-MS can substantially improve oligonucleotide detection sensitivity while reducing background and metal-adduct formation.

It should be noted that this comparison was made between self-packed capillary HILIC columns and commercial analytical HILIC columns. Further improvement on capillary column packing, stationary phase chemistry, frit preparation, and emitter configuration may further improve chromatographic performance and MS sensitivity. Nevertheless, the current capillary HILIC-MS setup still provided effective oligonucleotide separation while offering improved MS sensitivity and reduced background and adduct formation.

### Sample preparation strategy for low-input biological RNA samples

To enable robust and sensitive analysis of complex biological RNA samples with capillary-HILIC system, digested RNA fragments needed to be dissolved in a small volume of sample-loading buffer. Based on the optimization above, 70% ACN containing 10 mM ammonium acetate (AmAA) was selected as the loading buffer. However, initial attempts to dry silica spin column purified RNA, redissolve it in a 3 µL of water with 33 mM AmAA, and then add 7 µL acetonitrile resulted in visible white precipitates. This was likely caused by residual nonvolatile salts, such as Na⁺ and K⁺, carried over from the silica spin-column purification workflow and condensed into small volume after drying.

As shown in Figure 3, we addressed this limitation by adapting a C18 pipette-tip cleanup protocol commonly used for oligonucleotide preparation before MALDI–TOF analysis. In this workflow, RNA fragments are retained on the C18 material in the presence of an ion-pairing reagent, while salts and other impurities are removed during washing. Residual ion-pairing reagent is subsequently removed by washing with water, and the RNA fragments are eluted directly into the high-organic loading buffer used for capillary HILIC–MS. Using Agilent OMIX C18 10-µL pipette tips, up to 7 µg of digested RNA fragments could be purified and eluted in 6–10 µL of loading buffer for direct capillary HILIC–MS analysis. The approach was also adapted for analytical-scale sample preparation using Agilent OMIX C18 100-µL pipette tips, which enabled purification of up to 70 µg of digested RNA fragments. Under the standard workflow, 10 µg of input RNA was digested, yielding approximately 5–7 µg of RNA fragments after spin-column cleanup. This amount was compatible with a single 10-µL C18 pipette tip and was sufficient for triplicate LC–MS analyses. The method was also successfully applied to 3 µg of input RNA for duplicate analyses in the Human RNome benchmark study. In a proof-of-principle low-input experiment, 1 µg of RNA was digested and eluted in 4 µL of loading buffer and successfully analysed by injecting the entire sample. Overall, this workflow provides a practical and scalable sample-preparation strategy for LC–MS analysis of low-input biological RNA samples.

**Figure 3.**
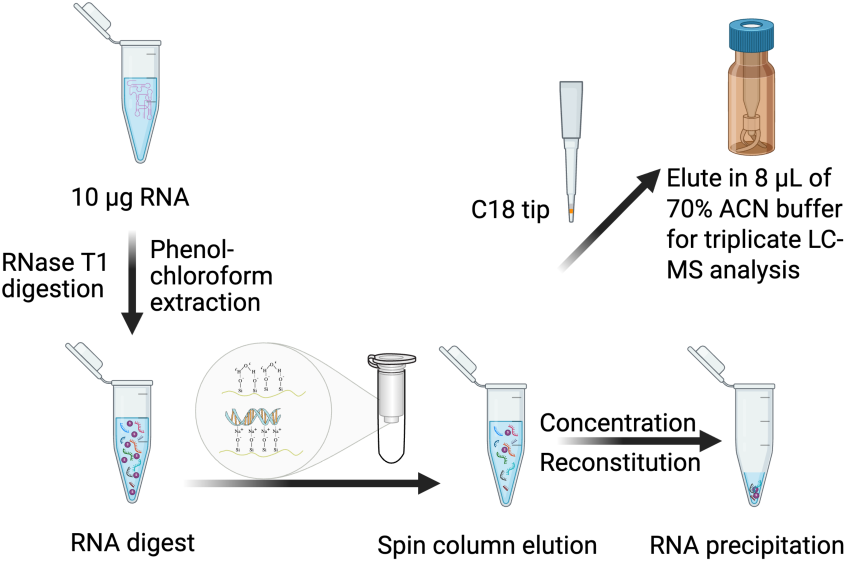
Schematic overview of the sample-preparation workflow used to desalt and concentrate RNA oligonucleotides with a C18 tip, followed by elution into a high-organic solvent compatible with HILIC–MS injection. The schematic was created with BioRender.

### Reproducible capillary HILIC-MS analysis of HEK293 rRNA digests

To assess analytical reproducibility, 10 µg of HEK293 rRNA was digested with RNase T1 and processed using the optimized sample-preparation workflow. Approximately 5 µg of RNA fragments were recovered after spin-column purification, desalted using a C18 peptide tip, and eluted in 8 µL of 70% ACN containing 10 mM AmAA. Three 2 µL technical replicate injections were analyzed by capillary HILIC–MS/MS. Raw data were processed using OpenMS, with Nucleic Acid Search Engine (NASE) for oligonucleotide identification and mapping. Mapped fragments passing an FDR threshold of ≤0.1 were retained for downstream analysis in Python. Retention time (RT) and peak intensity values were extracted from the featureXML files and grouped according to feature validity. Features annotated as “valid” were considered successfully fitted chromatographic features, whereas features with boundary-related fitting warnings were classified as non-valid. In the RT and peak-intensity reproducibility plots, teal points indicate features annotated as valid in both compared replicates, while orange points indicate features annotated as non-valid in at least one replicate.

Across the three replicate injections, the method showed broadly consistent chromatographic performance of complex rRNA digests. The total ion chromatograms were highly similar across the three injections (**Fig. 4a**), indicating stable overall separation behavior and comparable sample loading. RNA fragments generally eluted in order of increasing molecular mass, although sequence-dependent deviations were observed (**Fig. 4b**). These deviations likely reflected differences in nucleotide composition; for example, C-rich fragments tended to elute later than A-rich fragments. Retention-time reproducibility was high for features with valid peak models, with Pearson’s r values greater than 0.999 across replicate comparisons, although several outliers were observed (**Fig. 4c**). Most of these outliers were associated with non-valid feature-model statuses, particularly in the low-retention-time region. Further inspection suggested that these outliers may arise from interfering signals within the 5-ppm *m/z* extraction window, including partially overlapping features, nearby chromatographic peaks with different MS1 isotope distributions, or background signals, particularly in the low-retention-time region (**Fig. S4a**). Manual inspection revealed that the two outliers classified as valid features in the Rep1–Rep2 and Rep2–Rep3 comparisons arose from ambiguous peak assignment for the same precursor *m/z*. Although two chromatographic peaks were present in all three replicates, the software inconsistently selected the earlier-versus later-eluting peak across injections (**Fig. S4b**).

**Figure 4.**
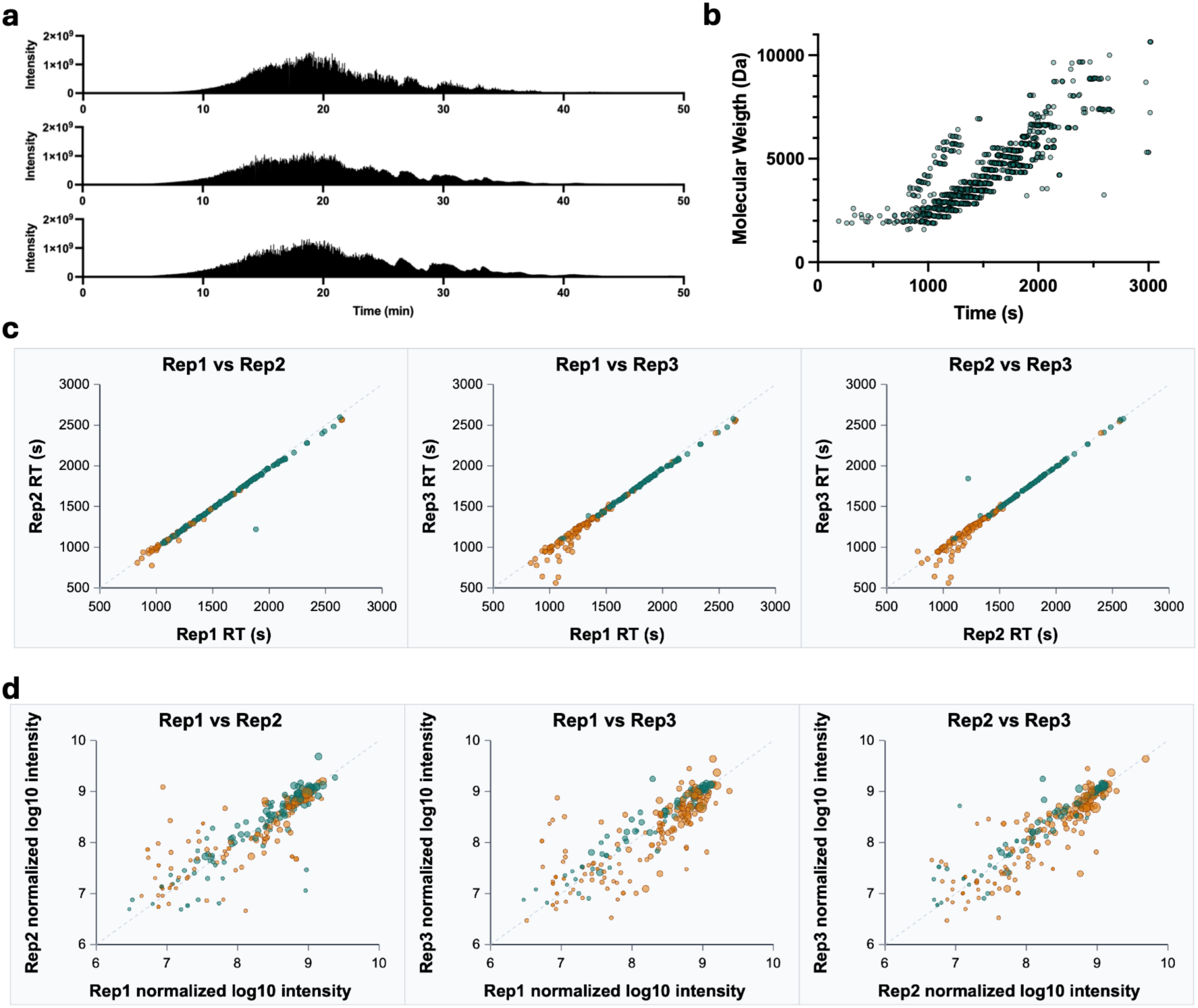
Reproducibility of capillary HILIC–MS analysis of digested HEK2G3 rRNAs. (**a**) TICs from three technical replicate injections. (**b**) Molecular weight and retention time of identified RNA fragments. (**c**) Pairwise retention-time comparisons of precursor features detected in all three injections and matched by *m/z* and charge state within 5 ppm. The diagonal line indicates perfect agreement. Teal points indicate features classified as valid in both injections, whereas orange points indicate features classified as non-valid in at least one injection. (**d)** Pairwise comparison of shared sequences detected in all three injections based on normalized log_10_ summed precursor intensities. Intensities were normalized to the total precursor intensity of each injection, with injection 1 used as the reference. Colors indicate feature validity as defined in (**c**).

Peak-intensity reproducibility was lower than retention-time reproducibility across replicate injections (**Fig. 4d**). As observed for RT reproducibility, feature validity strongly affected the intensity comparison: many features with non-valid peak-model statuses showed poor intensity agreement, consistent with inaccurate peak extraction or poorly defined chromatographic boundaries. However, even after excluding invalid features, intensity reproducibility remained lower than RT reproducibility, with Pearson’s r values of 0.889 for Rep1 vs Rep2, 0.944 for Rep1 vs Rep3, and 0.917 for Rep2 vs Rep3. This may reflect additional variability in nanospray stability from the capillary emitter, which can influence ionization efficiency and contribute to injection-to-injection differences in MS signal intensity. Together, these results indicate that retention time was highly stable across replicate injections, whereas peak intensity was more sensitive to emitter-dependent ionization variability.

### RNA modification maps of HEK293 rRNAs

We next evaluated the RNA modification maps of HEK293 rRNAs generated using NASE (**Figure S5**). The search was performed against *in silico* RNase T1-digested rRNA fragments, allowing a maximum of two variable modifications per fragment. One missed cleavage was permitted to enable detection of fragments containing Gm or m^7^G modifications, which can interfere with RNase T1 digestion. A total of 1,036 unique sequences were detected from 5,671 identified spectra that with q values ≤ 0.1. Venn diagram analysis showed that 229 fragments were detected in all three replicate injections, corresponding to 4,451 spectra, or 78% of the total identified spectra (Fig. 5a). This indicates that the majority of MS/MS spectra were assigned to reproducibly detected rRNA fragments, supporting the robustness of oligonucleotide identification across replicate capillary HILIC-MS injections.

**Figure 5.**
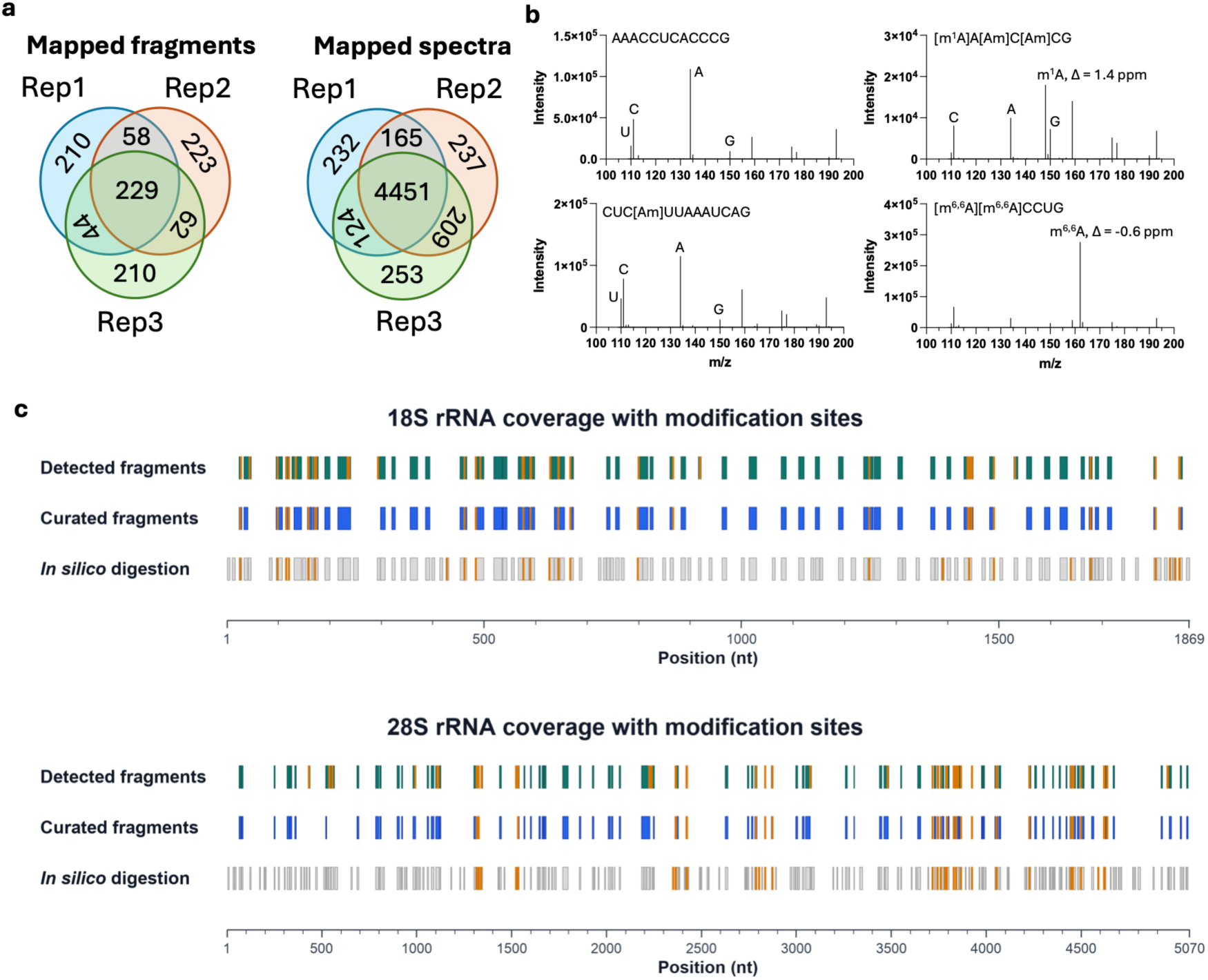
RNA modification mapping in HEK2G3 rRNAs. (**a**) Venn diagrams showing the numbers of mapped RNA fragments (left) and matched spectra (right) across three replicate injections. (**b**) Differentiation of Am, m^1^A, and m^6,6^A based on diagnostic nucleobase fragment ions. (**c**) Sequence-coverage maps of 18S (top) and 28S (bottom) rRNAs showing fragments detected across the three injections. Identified modification sites are indicated by orange markers.

We then analyzed these 229 fragments against previously reported rRNA modifications [42-48] and found that 131 fragments from 3,389 identified spectra were consistent with prior annotations (**Fig. 5c**). The remaining 98 unmatched fragments were then examined in more detail, as they highlight several current challenges in MS-based RNA modification analysis (**Supplementary Table 1**). First, 34 fragments were assigned to nearby positions within the rRNA sequence, typically differing by only one nucleotide. These cases likely reflect ambiguity in MS/MS ion-based modification mapping, where highly similar fragmentation patterns make it difficult to confidently determine the exact origin of the modification. Second, 24 fragments were assigned to modification isomers with same molecular weights, such as Am/m^6^A, Gm/m^7^G, and Cm/m^5^C. This reflects a common challenge in MS-based RNA modification analysis, where distinguishing base modifications from ribose methylations can be difficult and often depends on the presence or absence of diagnostic a–B ions. Third, 5 fragments were assigned to sequence isomers with identical precursor masses and highly similar sequences. Fourth, 6 fragments contained three methylation sites but were initially misassigned with m^6,6^A representing two methyl groups, because the NASE search was set to a maximum of two variable modifications per fragment. A subsequent search allowing up to three methylations resolved these assignments and confirmed the correct modification composition. Additionally, 10 fragments corresponded to unmodified or partially modified forms relative to the reference annotations. Six fragments containing unreported methylations appeared to arise from ion adducts of an unmodified precursor. The remaining 13 fragments could not be confidently assigned to a specific source and may represent false-positive identifications from the software-based search.

Based on these inspections, we removed incorrect modification identities and confirmed the most likely isomer assignments. The consensus results from the three replicate injections identified 56 modification sites, including 1 site in 5.8S rRNA, 24 sites in 18S rRNA, and 31 sites in 28S rRNA. These modifications comprised 17 Am, 1 m^1^A, 2 m^6,6^A, 8 Cm, 1 m5C, 15 Gm, 11 Um, and 1 m^1^acp^3^fi site. As we previously demonstrated that nucleobase fragments can be used for modification isomer identification in positive-ion mode, we found that this strategy can also distinguish Am, m^1^A, and m^6,6^A under negative-ion mode (**Fig. 5b**). In addition, we detected a low-abundance fragment carrying Um at the m^1^acp^3^fi1248 position. Based on the known biosynthetic pathway of m^1^acp^3^fi, we propose that this species most likely corresponds to the biosynthetic intermediate m^1^fi1248. This putative intermediate accounted for approximately 4% of the signal at this site, whereas the m^1^acp^3^fi modification represented the predominant species.

Together, these results show that most reproducibly detected spectra were consistent with known rRNA modification annotations, while the unmatched assignments largely reflected predictable limitations of MS/MS-based RNA modification mapping, including positional ambiguity, isomeric modifications, sequence isomers, search-space constraints, ion adducts, and potential false-positive assignments.

### Capillary HILIC-MS reveals rRNA modification heterogeneity

In addition to mapping annotated rRNA modifications, the improved sensitivity of the capillary HILIC-MS platform enabled simultaneous detection of several modified fragments together with their corresponding unmodified or partially modified counterparts. These paired detections provide an opportunity to estimate modification occupancy at specific rRNA regions, rather than simply recording the presence or absence of a modification. Using relative peak intensities, we observed incomplete modification at multiple sites, including 18S:Am99 (93%), 18S:Cm174 (87%), 18S:Am576 (62%), 18S:m^1^acp^3^fi1248 (96%), 18S:Cm1440 (11%), 18S:Um1442 (93%), 18S:Gm1447 (45%), 28S:Am1323 (49%), and 28S:Cm2365 (98%) (**Fig. 6**). Notably, 18S:Cm1440 showed particularly low modification occupancy in our dataset. Consistent with this low abundance, Cm1440 has primarily been reported by RiboMethSeq-based approaches [42,46] and has not been readily observed in X-ray crystal structures or previous MS-based rRNA modification analyses [43,48]. These results highlight the enhanced sensitivity of capillary HILIC-MS for detecting low-occupancy rRNA modifications and resolving co-existing modification states within complex rRNA populations.

**Figure 6.**
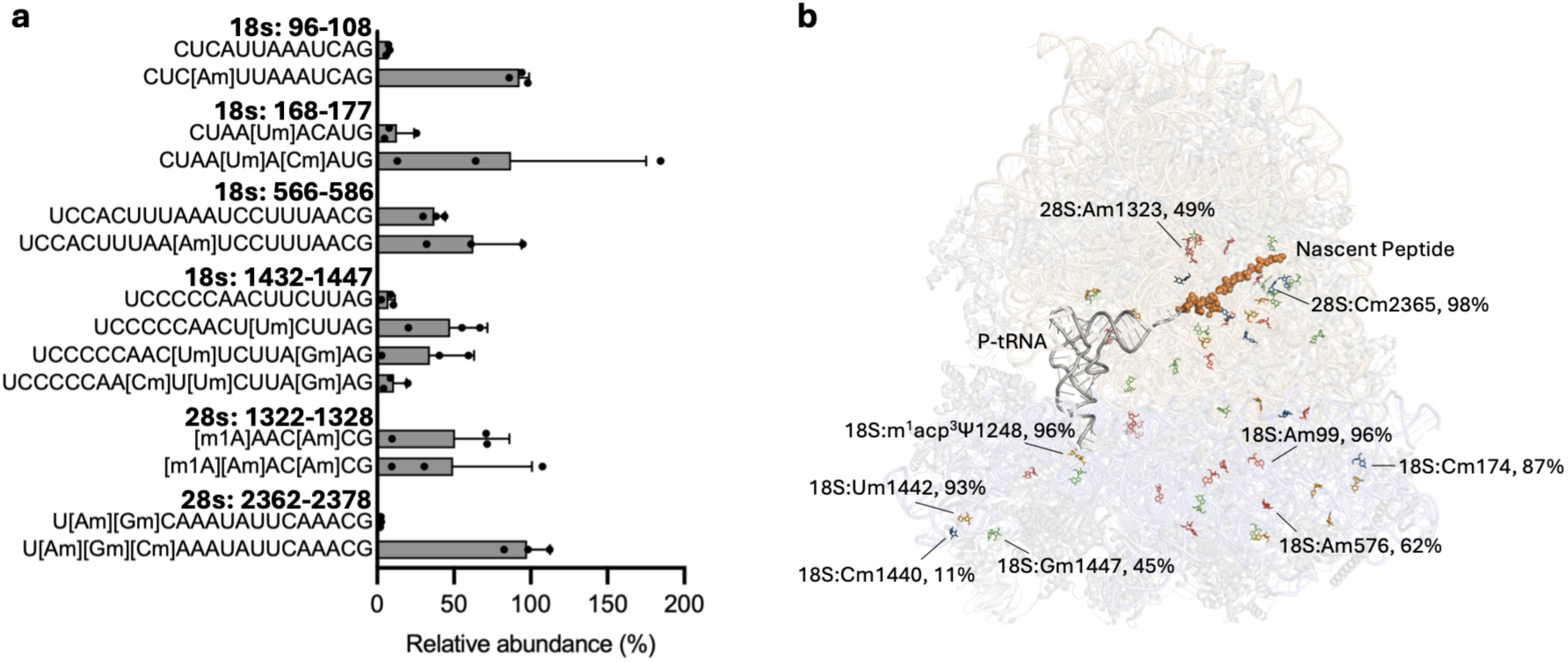
Partially modified sites in HEK2G3 rRNAs. (**a**) Bar plots showing six sets of RNA fragments detected in multiple modification states. The relative abundance of each state was calculated by normalizing its peak intensity to the total summed intensity of all states for the corresponding fragment. (**b**) Three-dimensional view of all mapped rRNA modifications in the human ribosome structure (PDB ID: 9O3V), together with the nascent peptide and P-site tRNA. Partially modified sites are labeled with their positions and estimated modification levels.

Many partial modification sites have previously been reported as variable or functionally responsive. For example, 18S:Cm174 has been reported as a dynamic MYC-responsive rRNA 2′-O-methylation site. It modulates translation of specific mRNA subsets, likely through effects on codon-dependent elongation and helix 8 dynamics in the small ribosomal subunit [49]. In a RiboMeth-Seq analysis of 195 primary human breast tumors, 18S:Am576 and 18S:Gm1447 were identified as variable rRNA 2′-O-methylation sites whose levels were significantly associated with breast cancer subtype, hormone receptor status, and tumor grade [50]. Specifically, 18S:Gm1447 methylation was decreased in triple-negative breast cancers compared with luminal tumors, whereas 18S:Am576 methylation was increased in triple-negative breast cancers and further distinguished benign from malignant mammary tumors. Similarly, 18S:Cm1440 and 18S:Gm1447 have been reported among hypomethylated in human cell and cancer-related studies [46,51]. Furthermore, hypomethylation of 28S:Am1323 has been reported in an RNA-binding protein FUS knockout model [52].

Together, these results indicate that capillary HILIC-MS can capture not only the presence of modified rRNA fragments, but also co-existing modification states at selected sites. Although differences in ionization efficiency, chromatographic behavior, and MS/MS sampling mean that paired modified/unmodified fragment signals should be interpreted as semi-quantitative rather than absolute stoichiometric measurements, they provide useful evidence for site-specific rRNA modification heterogeneity. This highlights the potential of capillary HILIC-MS as a complementary approach for studying dynamic modified rRNA sites in disease model.

### RNA modification mapping in PA14 total tRNAs

Modification mapping in total tRNA is more challenging than in rRNA because the greater complexity of tRNA modifications complicates data analysis and requires higher-quality MS/MS spectra for confident sequence assignment. To evaluate the performance of the capillary HILIC–MS platform for tRNA analysis, 10 µg of total tRNA isolated from *Pseudomonas aeruginosa* UCBPP-PA14 was processed using the same sample-preparation workflow as the rRNA samples. Retention-time comparisons demonstrated excellent reproducibility across the three technical replicate injections, with Pearson correlation coefficients exceeding 0.998 for all pairwise comparisons (**Fig. S6**). Modification mapping using NASE, with all detected modifications except inosine and pseudouridine specified as variable modifications, identified 663 unique sequences from 2,703 mapped spectra (**Fig. S6**). Of these, 454 sequences were detected in only one injection and were therefore considered more likely to represent false-positive assignments. In contrast, 80 sequences were reproducibly detected in all three injections, collectively accounting for 1,628 spectra, with a median of 13 spectra per sequence. These reproducibly detected sequences provided 14% tRNA sequence coverage. For comparison, *in silico* RNase T1 digestion yielded a theoretical coverage of 35% when only fragments ≥6 nt were considered. These results indicate that sequences repeatedly detected within and across injections provide stronger evidence for reproducible capillary HILIC-MS performance.

A total of 22 distinct modifications were mapped to 68 sites in PA14 tRNAs (**Fig. 7**). However, assignments of some isomeric modifications may remain uncertain, even though the ambiguous-assignment option in NASE was enabled to identify diagnostic a-B ions. Among these, four distinct wobble-position modifications were identified: Q, mnm^5^s^2^U, cmo^5^U and Cm (**Supplementary Table 2**). Q was observed in tRNA^Asn^_GTT_ and tRNA^Tyr^_GTA_. Both tRNA contain a single G34 anticodon that is used to decode both C- and U-ending codons. The conversion of G34 to Q34 enhances decoding of U-ending codons while maintaining recognition of C-ending codons, thereby expanding the decoding capacity of a single tRNA species. mnm^5^s^2^U was observed in tRNA^Lys^_TTT_; cmo^5^U was observed in tRNA^Val^_TAC_. These tRNAs use a single U34 anticodon to decode both A and G ending codons. The mnm^5^s^2^U and cmo^5^U modifications enhance wobble decoding by promoting accurate recognition of both codons while minimizing translational errors. Furthermore, multiple modification states were observed at position 37. Both io^6^A and ms^2^io^6^A were detected in tRNA^Cys^_GCA_ and tRNA^Ser^_CGA_, whereas ms^2^i^6^A and ms^2^io^6^A were detected in tRNA^Phe^_GAA_, tRNA^Trp^_CCA_ and tRNA^Tyr^_GTA_. Together, these results demonstrate that the capillary HILIC–MS workflow enables reproducible modification mapping in complex total tRNA samples, despite the greater analytical challenges associated with tRNA heterogeneity and modification density. Importantly, the ability to detect a broad range of chemically distinct modifications, including multiple modification states at the same nucleotide position, represents a key advantage of the MS-based platform. This capability provides a direct view of tRNA modification diversity that is difficult to obtain using sequencing-based approaches alone.

**Figure 7.**
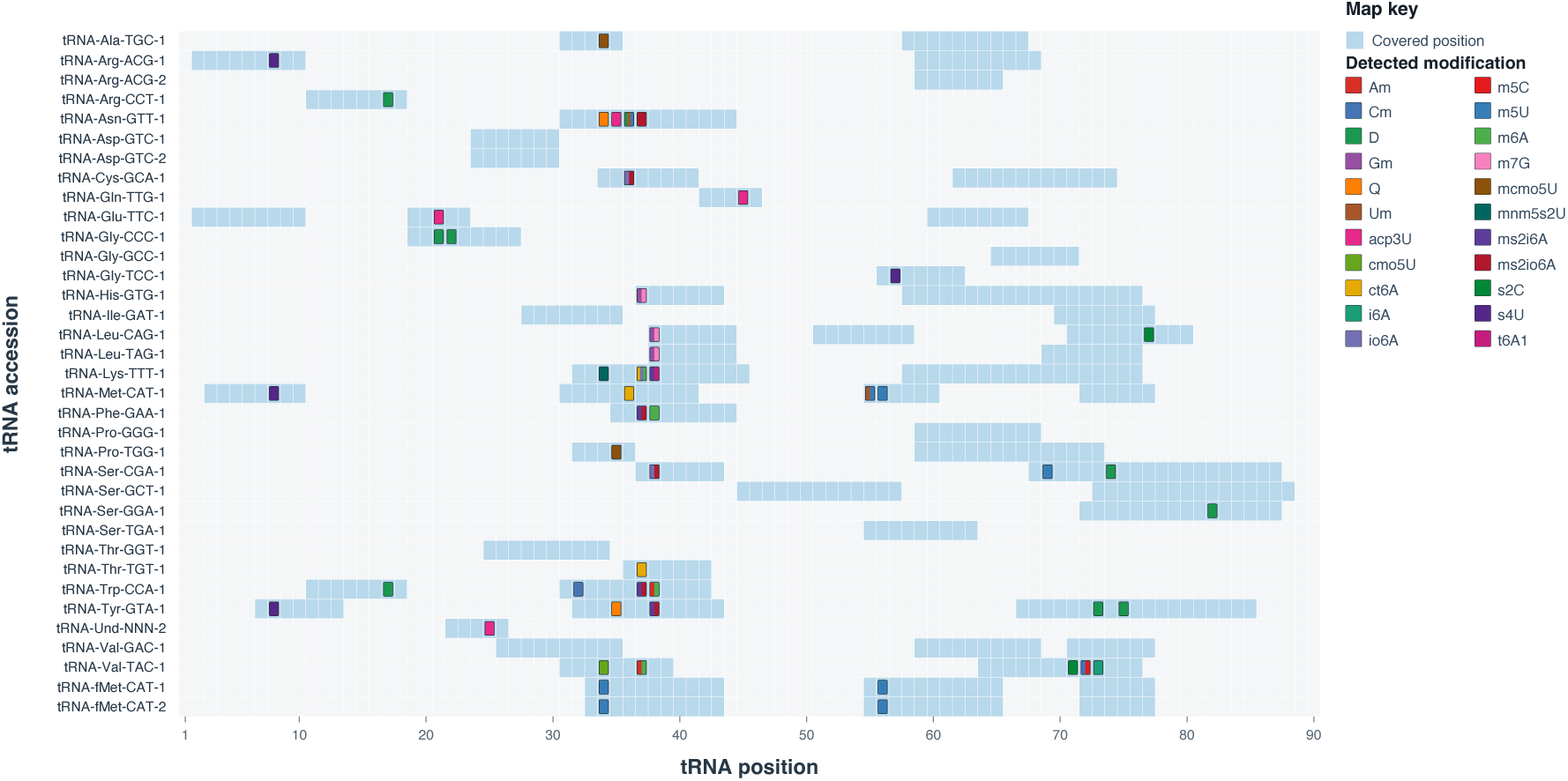
Modification map of 35 PA14 tRNAs. All reported assignments had a false discovery rate of *q* ≤ 0.1 and were detected in all three replicate injections. Detected sequence positions are shown in light blue, and modified nucleotides are indicated using the colors defined in the legend. For PA14 tRNAs in which the anticodon is annotated at positions other than the conventional 34–36, the original positional numbering was retained without adjustment.

## CONCLUSION

In conclusion, we established an ion-pair-free capillary HILIC–MS workflow that combines a metal-free capillary configuration, high-organic sample loading, optimized volatile-buffer and source conditions, and C18 tip-based cleanup for sensitive analysis of oligonucleotides and digested biological RNAs. Compared with analytical-scale HILIC–MS, the capillary platform substantially improved MS sensitivity while reducing background signals and metal-adduct formation. It also provided highly reproducible chromatographic retention across technical replicate injections, although peak-intensity reproducibility remained more sensitive to nanospray stability.

Application of the workflow to HEK293 rRNAs and PA14 tRNAs identified 56 and 68 modification sites, respectively. The results demonstrated that negative-mode diagnostic nucleobase ions can help distinguish selected isomeric modifications, including Am, m^1^A, and m^6,6^A. Detailed examination of assignments inconsistent with previously reported modification sites highlighted persistent challenges in oligonucleotide MS, including positional ambiguity, sequence and modification isomers, adduct-related misassignments, and other unidentified false positives. Although individual injections yielded a relatively large number of potentially incorrect sequence assignments, sequences reproducibly detected across all three injections were strongly enriched for high-confidence matches. These findings demonstrate the value of replicate-based filtering, together with manual spectral validation, improved search algorithms, and orthogonal confirmation, for the reliable analysis of complex RNA samples.

Importantly, the enhanced sensitivity enabled the simultaneous detection of multiple modified, and partially modified site states, providing semi-quantitative evidence of site-specific RNA modification heterogeneity. These included 9 partially modified sites in rRNAs, several of which have been linked to disease-related processes, the low-occupancy 18S:Cm1440, which is difficult to detect using structural methods or conventional MS approaches. Multiple modification states were also observed at position 37 of several tRNAs. However, differences in chromatographic behavior, and ionization efficiency can affect relative signals, these measurements should not yet be interpreted as absolute modification stoichiometries. More accurate quantification strategies, including label-free approach using internal standards and isobaric mass-tagging methods compatible with negative-ion MS, would be valuable for future studies.

The absence of ion-pairing reagents also facilitates switching the LC–MS system back to proteomics applications without extensive system cleaning or source decontamination. Returning the Easy-nLC 1000 to the proteomics workflow required approximately 2–3 hours for solvent purging and calibration. This practical compatibility allows a high-end LC–MS platform to be shared between RNA and proteomics workflows with substantially less disruption than workflows relying on persistent ion-pairing reagents.

Overall, this workflow provides a practical and accessible approach for low-input RNA modification analysis on high-performance LC–MS platforms. Further improvements in capillary-column manufacturing, emitter stability, quantitative methods, and computational identification should expand its utility for characterizing the chemical diversity, heterogeneity, and dynamics of RNA modifications in biological and disease-related samples.

## ACKNOWLEDGEMENTS

We thank Jordy Hsiao and Connor Flannery from Agilent Technologies for valuable advice on HILIC method development and troubleshooting.

## AUTHOR CONTRIBUTIONS

Junzhou Wu: Conceptualization, Investigation, Formal analysis, Methodology, Visualization, Project administration, Supervision, Writing—original draft, Writing – review & editing. Rana Togay: Investigation, Formal analysis, Methodology, Writing— original draft. Jingjing Sun: Investigation, Methodology, Writing—review & editing. Dwijapriya: Investigation, Resources, Writing—review & editing Chi-Kong Chan: Investigation, Writing—review & editing. Alex Reading: Investigation, Writing—review & editing. Liang Cui: Resources, Writing—review & editing. Xueming Dong: Funding acquisition, Supervision, Writing—review & editing. Peter Dedon: Conceptualization, Funding acquisition, Project administration, Supervision, Writing—review & editing.

