## Supplementary Data for "Ion-Pair-Free Capillary HILIC-MS for Sensitive Nucleic Acid Analysis and RNA Modification Mapping"

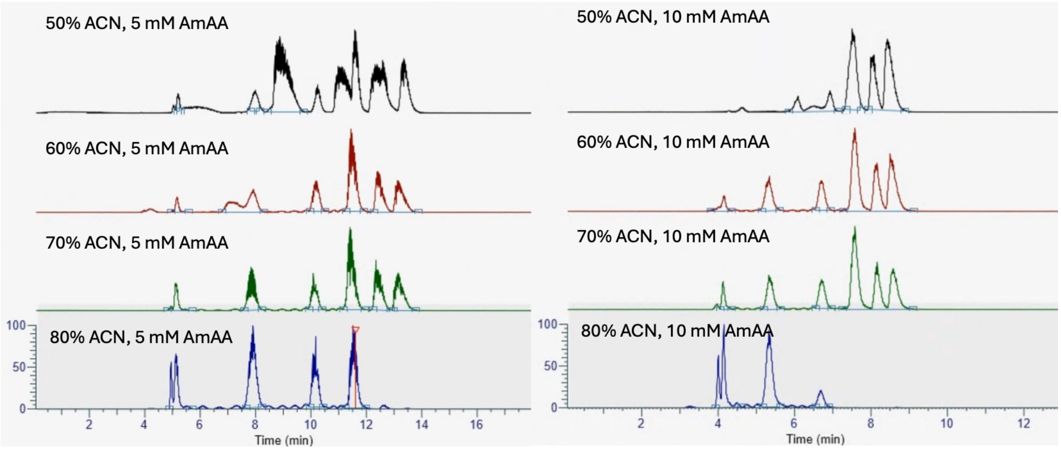


**Figure S1**. Comparison of total ion chromatograms (TICs) of 5–30 nt poly(T) oligonucleotides using loading buffers containing 50–80% acetonitrile and 5 or 10 mM ammonium acetate, analyzed by capillary HILIC–MS.

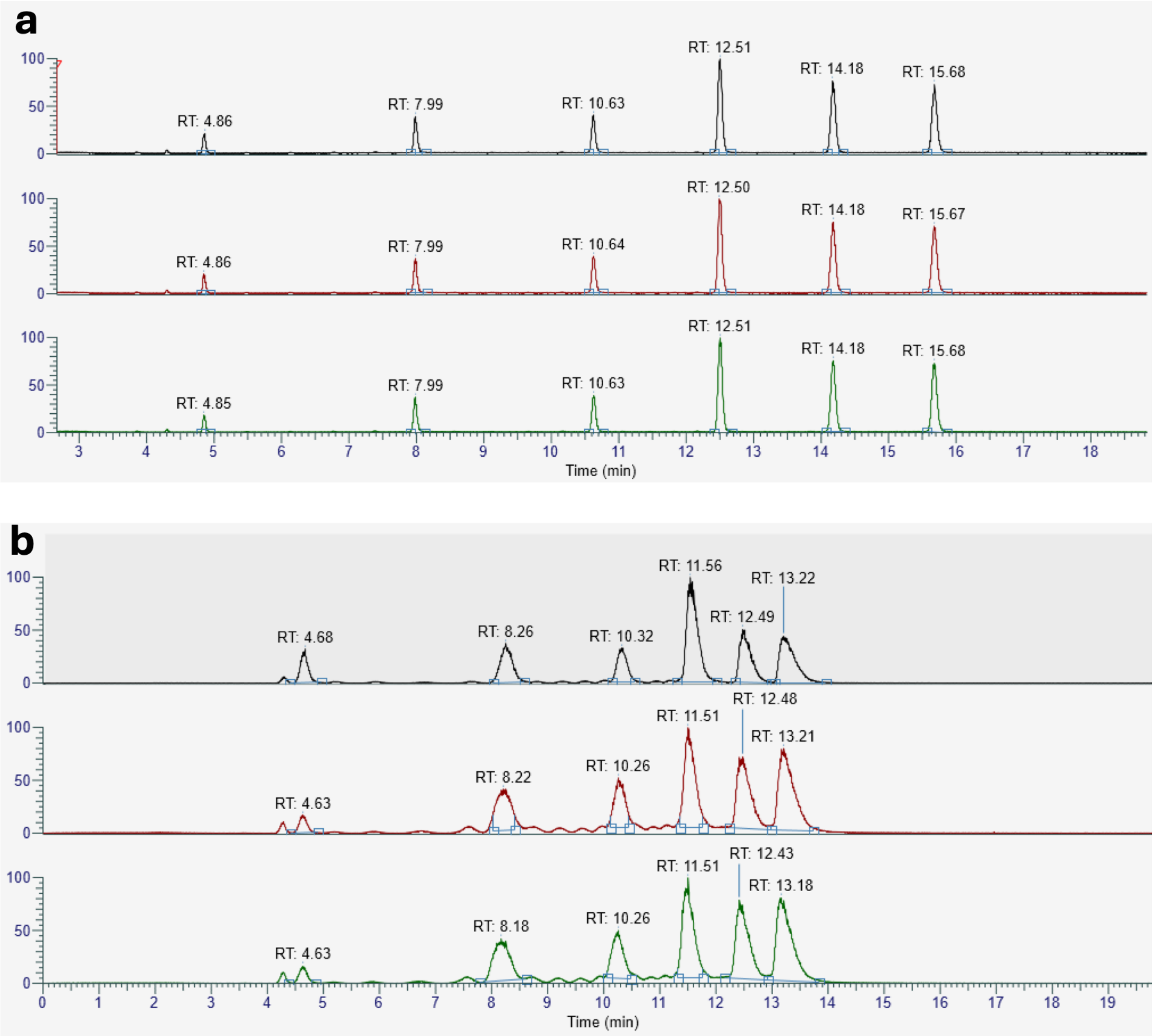


**Figure S2**. Triplicate injections of 5–30 nt poly(T) oligonucleotides analyzed using an analytical HILIC column (**a**) and a capillary HILIC column (**b**), shown as TICs.

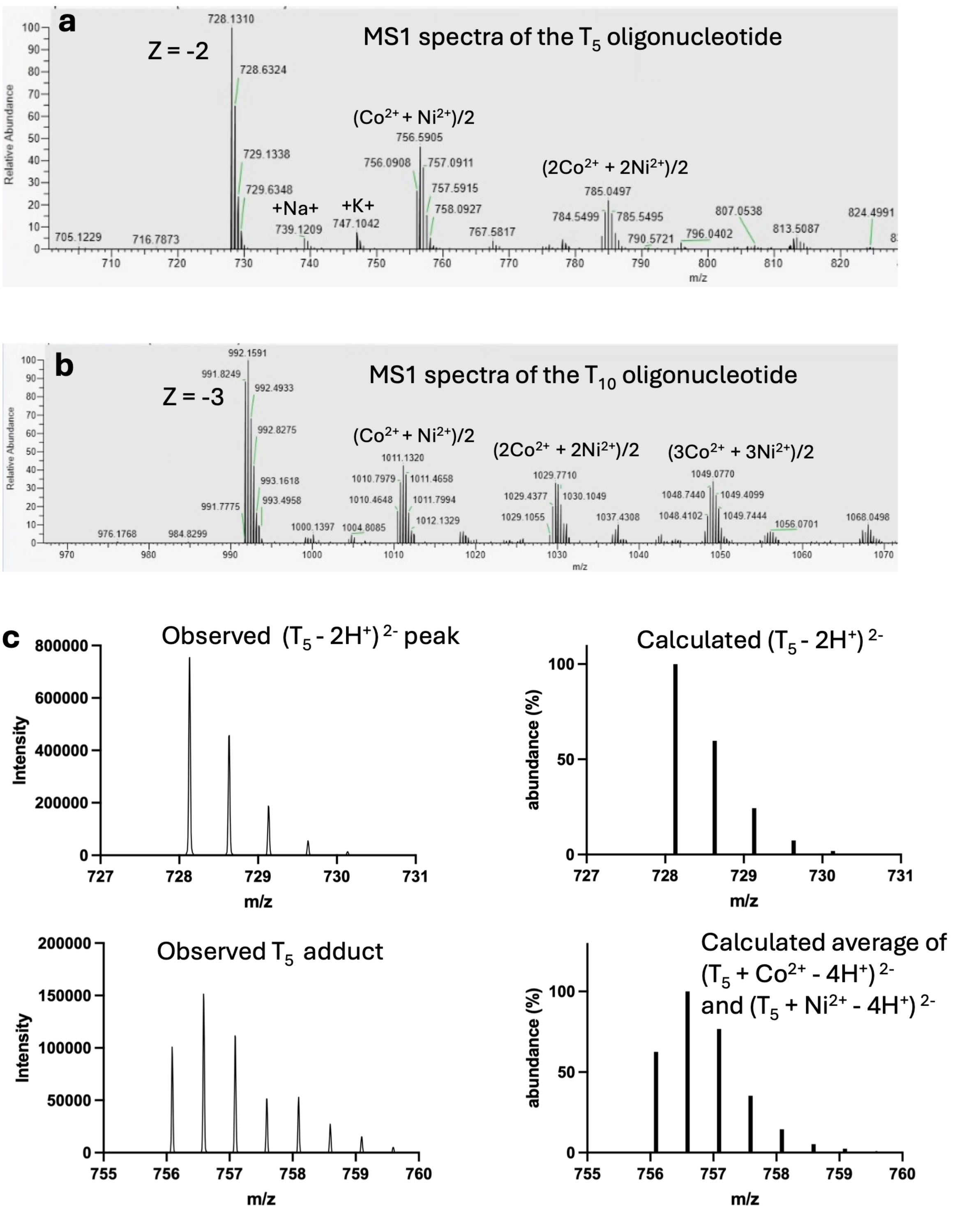


**Figure S3**. Representative MS1 spectra of 5-nt (**a**) and 10-nt (**b**) oligonucleotides showing a series of ion adducts. The major abnormal adduct peaks exhibited isotopic distributions inconsistent with conventional sodium or potassium adducts. (**c**) Enlarged view comparing the observed isotopic distributions of the molecular ion and abnormal adduct ion with the calculated distributions for the molecular ion and a mixed $(Co^{2+} + Ni^{2+})/2$ adduct.

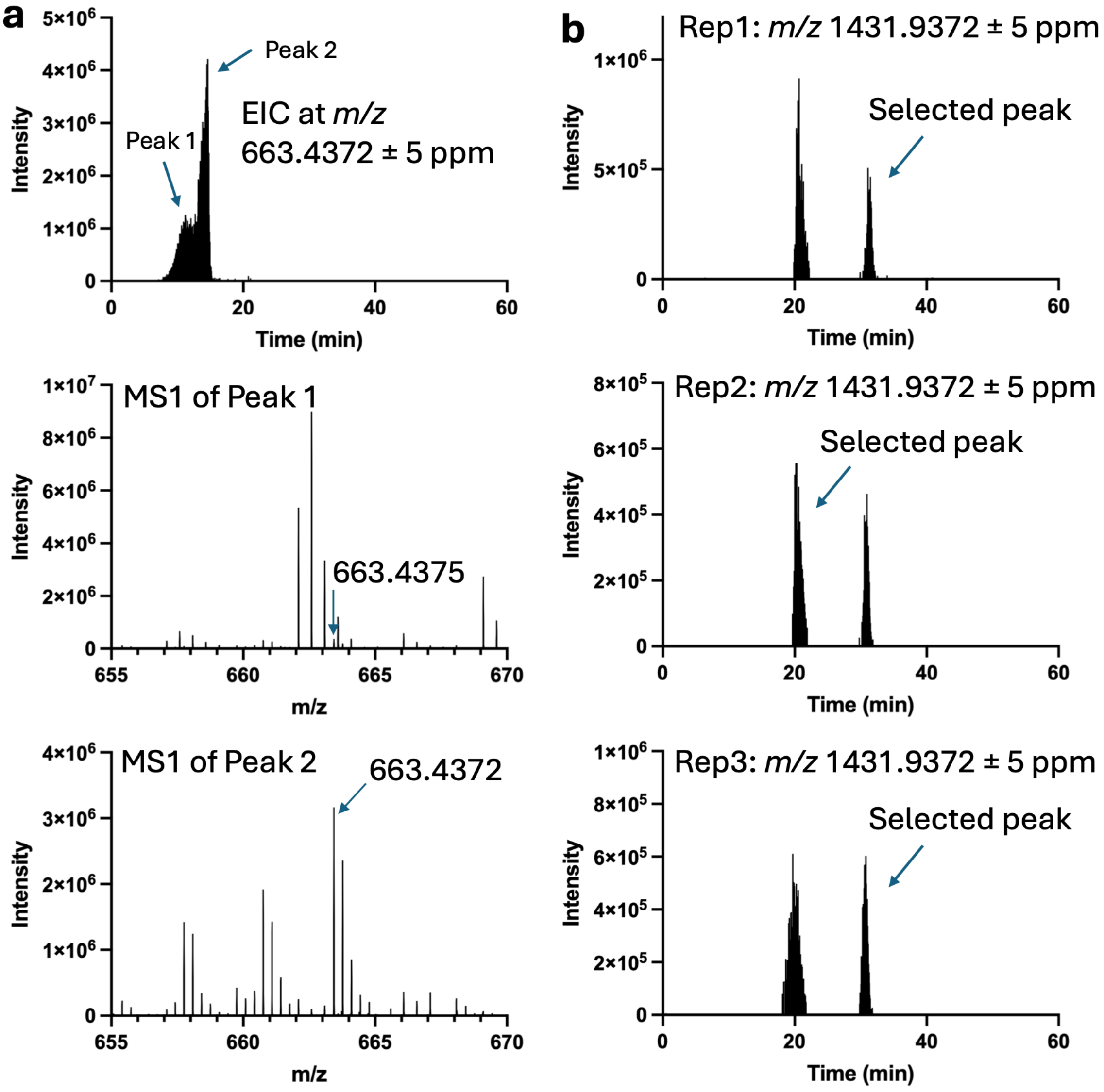


**Figure S4. Inspection of outlier features across replicate comparisons.** **(a)** Representative outlier associated with a non-valid feature-model status. Top: Extracted ion chromatogram (EIC) of *m/z* 663.4372 using a 5-ppm extraction window, showing two partially merged chromatographic peaks. Middle: MS1 spectrum of peak 1, showing an ion at *m/z* 663.4375. Bottom: MS1 spectrum of peak 2, showing an ion at *m/z* 663.4372. **(b)** EICs of m/z 1431.9372 using a 5-ppm extraction window for Rep1 (top), Rep2 (middle), and Rep3 (bottom). Arrows indicate the peaks selected by the OpenMS pipeline for label-free quantification.

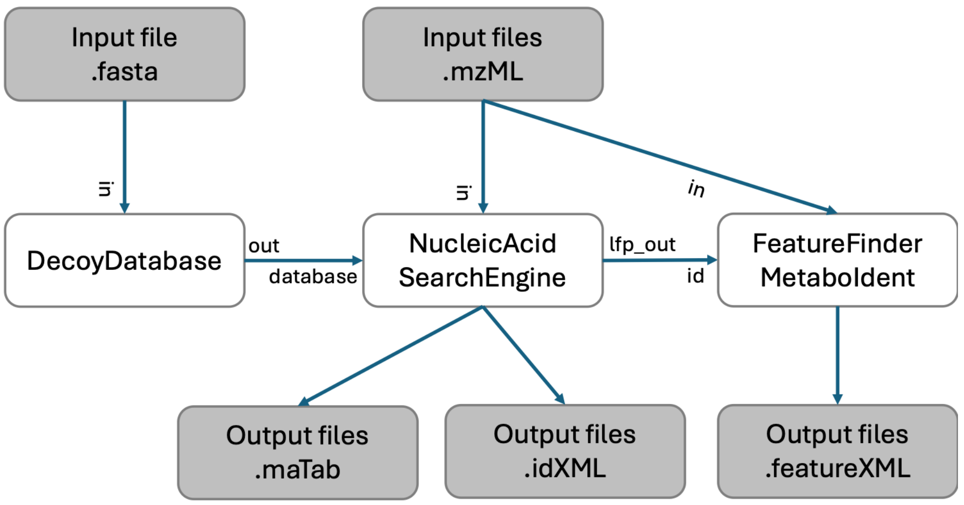


**Figure S5**. OpenMS workflow for MS-based RNA modification mapping and label-free quantification.

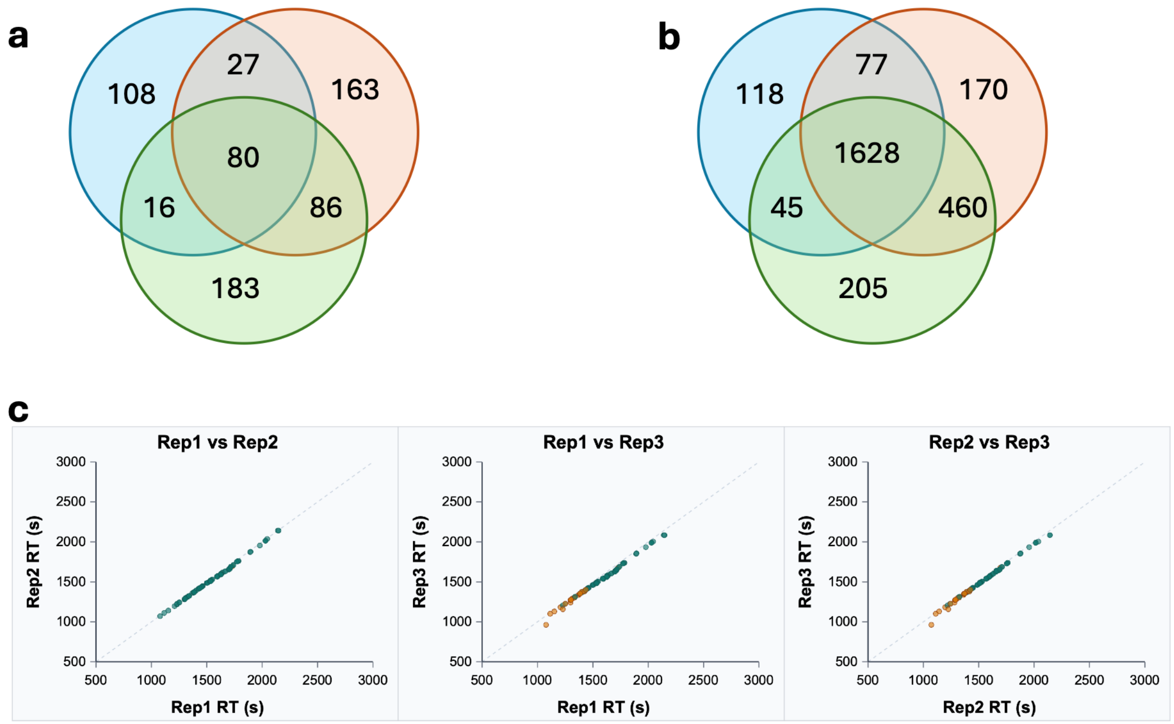


**Figure S6. RNA modification analysis in PA14 tRNAs.** Venn diagrams showing the numbers of mapped RNA fragments (**a**) and matched spectra (**b**) across three replicate injections. (**c**) Pairwise retention-time comparisons of precursor features detected in all three injections and matched by *m/z* and charge state within 5 ppm. The diagonal line indicates perfect agreement. Teal points indicate features classified as valid in both injections, whereas orange points indicate features classified as non-valid in at least one injection.
